# Screening of anti-mycobacterial compounds in a naturally infected zebrafish embryo model

**DOI:** 10.1101/038810

**Authors:** J.P. Dalton, B. Uy, K.S. Okuda, C.J. Hall, W.A. Denny, P.S. Crosier, S. Swift, S. Wiles

## Synopsis

*Mycobacterium tuberculosis* is a deadly human pathogen that latently infects a third of the world's population, resulting in approximately 1.5 million deaths per year. Due to the difficulties and expense of carrying out animal drug trials using *M. tuberculosis* and rodents, infections of the zebrafish *Danio rerio* with *M. marinum* have been used as a surrogate. However the methods so far described require specialised equipment and a high level of operator expertise.

We investigated a natural infection model where zebrafish embryos are infected through incubation in media containing *M. marinum*. Using bioluminescently labelled *M. marinum*, we have characterised the nature of infection and established a model for interventional drug therapy. We have used a selection of traditional and experimental compounds to validate this model for anti-mycobacterial drug discovery. We observed that only three of the six treatments tested (Delamonid, SN30527 and rifampicin) retarded the growth of *M. marinum* in vitro. In contrast, five of the six treatments (Pretomanid, Delamanid, SN30488, SN30527 and rifampicin) retarded the growth of *M. tuberculosis* in vitro. Importantly, these same five treatments significantly reduced the bioluminescent signal from naturally infected zebrafish embryos.

Overall this study has demonstrated that zebrafish embryos naturally infected with bioluminescent *M. marinum* M can be used for the rapid screening of anti-mycobacterial compounds with readily available equipment and limited expertise. The result is an assay that can be carried out by a wide variety of laboratories for minimal cost and without high levels of zebrafish expertise.

## Introduction

*Mycobacterium tuberculosis* is a deadly human pathogen that latently infects a third of the world's population. Around 5-10% of these infections develop into active disease, resulting in approximately 1.5 million deaths per year^1^. Tuberculosis (TB) is treatable with antibiotics, albeit with extended treatment times required and a high financial cost^1^, and as a result has become less prevalent in the developed world, although it remains a major health issue globally^1^. However the emergence of multi drug resistant (MDR-TB) and totally drug resistant (TDR-TB) isolates of *M. tuberculosis* has once again brought TB into the spotlight for all countries and has necessitated the development of novel drugs to combat this pathogen ^2, 3^. A change in the global disease burden of HIV/AIDS has also added impetus to this cause; the number of people co-infected with HIV and *M. tuberculosis* is rapidly increasing globally, resulting in a destructive synergy which exponentially exacerbates the disease progression of both diseases ^4^.

Due to the difficulties and dangers involved in culturing *M. tuberculosis*, an airborne Biosafety Level 3 pathogen, faster-growing and less pathogenic mycobacterial species, such as *M. smegmatis* and *M. marinum*, are routinely exploited for TB research and anti-mycobacterial drug discovery^5–8^. *M. marinum* is a pathogen of ectotherms (fish, amphibians and reptiles) that produces a tuberculosis-like disease ^9^. *M. marinum* is a close genetic relative of *M. tuberculosis*, with which it shares conserved virulence determinants ^9^, and has been known to cause granulomatous skin infections in humans ^10, 11^. Infection of the tropical zebrafish, *Danio rerio*, with *M. marinum* has been used to develop a surrogate in vivo model of TB pathogenesis ^12^ useful for the rapid screening of potential antimycobacterial compounds^13^. Zebrafish are genetically tractable ^9^ and possess a complex immune system comparable to that of humans ^14–16^. As a result, zebrafish have been extensively used for disease modelling and drug discovery for both communicable and non-communicable diseases^13, 17–24^. Infection of zebrafish with *M. marinum* through microinjection results in the development of necrotic granulomatous lesions reminiscent of human TB infection,^25, 26^ lending weight to the use of this model host as a surrogate for mycobacterial research and drug discovery. One drawback of the previously published infection protocols is that they require specialised equipment and a high level of operator expertise^13^.

Bioluminescence is a biological reaction which results in the production of light via a luciferase catalysed reaction. It is a naturally occurring process with several variants seen across several kingdoms and has been harnessed as a reporter in both in vitro and in vivo assays^27, 28^Bioluminescence allows for non-invasive monitoring of luciferase-expressing bacteria within a host, as the light produced by the bacterium travels through the host tissues and can be readily detected ^28–30^. As tagged cells only produce a signal when alive, bioluminescence is an excellent reporter to rapidly assay for antimicrobial compounds, non-destructively and in real-time, in microtitre plate formats using a luminometer, or in vivo using sensitive imaging equipment^28, 31–33^.

Zebrafish and other fish readily become naturally infected with *M. marinum* present in their environment^34^. However it is not possible to tell which fish are infected at an early stage without euthanising the animal and plating out for viable bacteria^26^. Here we use bioluminescently tagged *M. marinum* M to establish natural infections in zebrafish embryos, utilising the light emitted by these bacteria to identify which fish have become infected. We show that naturally infected fish can be treated with potential anti-mycobacterial compounds and light output used as an indicator of in vivo drug efficacy.

## Materials and Methods

### Bacterial strains and plasmids

Bacterial strains and plasmids used throughout this study are described in Table 1. Bacteria were transformed as previously described ^35^. *M. marinum* strains were grown at 28°C and *M. tuberculosis* at 37°C, with shaking at 200rpm and 100rpm respectively, in Middlebrook 7H9 broth (Fort Richard) supplemented with 10% ADC enrichment media (Fort Richard) and 0.5% glycerol (Sigma Aldrich) under the appropriate antibiotic selection (kanamycin at 25μg ml^-1^ and hygromycin at 50μg ml^-1^[Sigma Aldrich]).

**Table 1.** Strains and plasmids used in this study.

| Strain/plasmid | Description | Reference |
| --- | --- | --- |
| *M. marinum* ATCC BAA-535 | Wildtype isolate M |  |
| *M. marinum* BSG100 | ATCC BAA-535 containing pMV306G13FFlucRT chromosomally integrated, Km resistant. | This study |
| *M. marinum* BSG101 | ATCC BAA-535 containing pMV306G13LuxABCDE chromosomally integrated, Km resistant. | This study |
| *M. marinum* BSG102 | ATCC BAA-535 expressing pTEC27, Hyg resistant. | This study |
| *M. marinum* BSG103 | BSG101 expressing pTEC27, Km and Hyg resistant. | This study |
| *M. tuberculosis* BSG001 | *M. tuberculosis* ATCC (H37Rv) containing MV306hsp + LuxAB + G13 + CDE chromosomally integrated, Km resistant. | 50 |
| pMV306G13FFlucRT | Mycobacterial integrating vector containing a modified firefly luciferase gene optimised for use in Mycobacteria, Km resistant. | 49 |
| pMV306G13LuxABCDE | Mycobacterial integrating vector containing the *lux* operon from *Photobacterium luminescens* optimised for use in Mycobacteria, Km resistant. | 49 |
| pTEC27 | Mycobacterial plasmid containing the red fluorescent protein tdTomato, hyg resistant. | 13 |

Key: Km, kanamycin; hyg, hygromycin.

### In vitro drug testing

*M. marinum* BSG101 and *M. tuberculosis* BSG001 were grown without antibiotic selection to mid log phase, then diluted to an optical density at 600nm (OD_600_) of 0.01. Bacteria were aliquoted (100μl, 5x10^5^cfu ml^-1^ approx.) into the wells of a black 96 well microtitre plate (Grenier Bio-One). Test compounds were dissolved in DMSO, added 1:100 to the top most well and then diluted two-fold in a series down the plate (Table 2). The plates were incubated at 28°C *(M. marinum)* or 37°C (M. *tuberculosis)* and bioluminescence monitored daily over 7 days using a Victor X1 luminometer (Perkin Elmer).

**Table 2.**
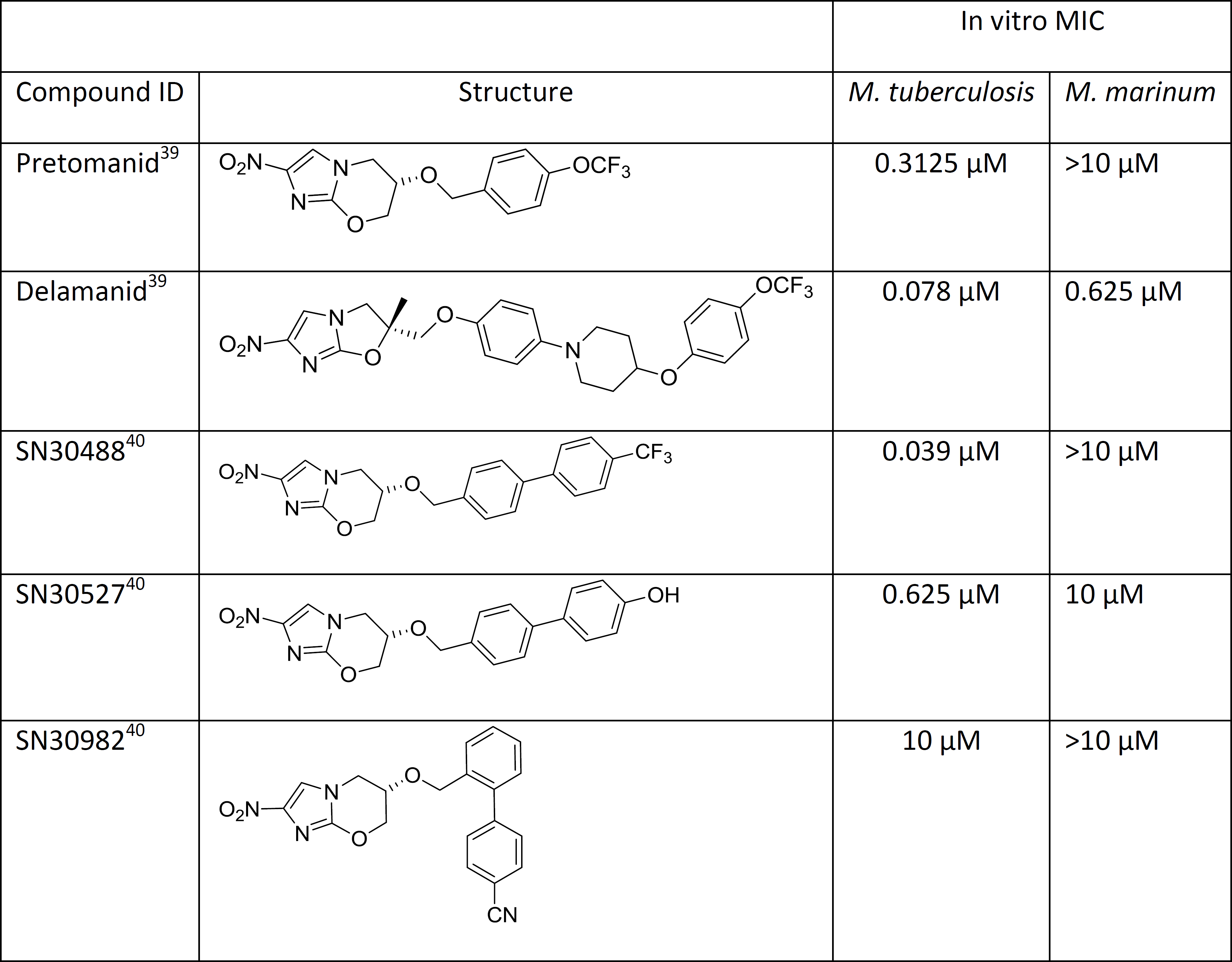
Novel compounds used in drug treatment assays and in vitro MIC values for *M. tuberculosis* BSG001 and *M. marinum* BSG101.

|  |  | In vitro MIC |  |
| --- | --- | --- | --- |
| Compound ID | Structure | *M. tuberculosis* | *M. marinum* |
| Pretomanid<sup>39</sup> |  | 0.3125 µM | >10 µM |
| Delamanid<sup>39</sup> |  | 0.078 µM | 0.625 µM |
| SN30488<sup>40</sup> |  | 0.039 µM | >10 µM |
| SN30527<sup>40</sup> |  | 0.625 µM | 10 µM |
| SN30982<sup>40</sup> |  | 10 µM | >10 µM |

Key: MIC, minimum inhibitory concentration.

### Fish husbandry

Zebrafish *(Danio rerio)* embryos were obtained from natural spawnings and raised at 28°C in E3 Medium (0.33 mM calcium chloride, 0.33 mM magnesium sulphate, 0.14mM potassium chloride and 5 mM sodum chloride). The medium was supplemented with 0.003% phenylthiourea (PTU) to inhibit pigmentation when embryos were being imaged. Zebrafish embryos of the age used do not fall under the New Zealand Animal Welfare Act 1999 and so experiments did not require approval from an animal ethics committee.

### Natural infection of zebrafish embryos

*M. marinum* strains were grown to mid log phase (OD_600_ between 0.8 and 2) without antibiotic selection, washed once in E3 before being adjusted to an OD_600_ of 1. The approximate concentration should be around 5x10^8^ cfu ml^-1^; this was confirmed by retrospectively plating inocula onto 7H11 supplemented with 10% OADC and 0.5% glycerol. Zebrafish embryos at 2 days post-fertilisation (dpf) were dechorionated, either manually or using pronase, as previously described^36^. Groups of embryos (50-300) were then placed in 9mm petri dishes containing 25ml of E3 supplemented with varying concentrations of *M. marinum* (by varying the volume of adjusted bacteria) and incubated for 4 days at 28°C. After infection, any non-internalised bacteria were removed by gently washing embryos four times with fresh E3 in groups of 10 in separate wells of a 24 well tissue culture plates (BD Falcon) using gentle aspiration of the media. Embryos were left in the 24 well plate overnight to remove transient bacteria and then rinsed 4 times in E3 to remove the transient population. Embryos were then individually placed into the wells of a clear-bottomed black 96 well microtitre plate (Nunc) with 100μl of E3 and prepared for drug treatment.

### Injection of zebrafish embryos

*M. marinum* strains were prepared as above. Zebrafish embryos 2dpf were manually de-chorionated, anaesthetised using 0.168 mg ml^-1^ tricaine (Sigma-Aldrich) in E3 medium^37^ and infected by microinjection into the caudal vein as previously described^38^.

### Measurement of bioluminescence from zebrafish embryos

All bioluminescent measurements were carried out using a Victor X1 luminometer (Perkin Elmer) with a 1 second exposure time on an open filter. Bioluminescence from zebrafish infected with *M. marinum* BSG100 was visualised after addition of luciferin (30 μg ml^-1^) (Gold Biotechnology).

### Drug testing in zebrafish embryos

Infected zebrafish embryo were read on a luminometer after removal of transient bacteria and detectably infected embryos (RLU >20) were randomly distributed within drug treatment groups. Drugs were made up to 100x working concentration in an appropriate solvent. They were diluted 1:10 in E3 and 10μl of diluted drug was placed into each well of the 96 well plate containing 100μl of E3. The final concentration of the test compounds was 10μM. E3 was used as a no treatment control. Embryos were treated at 5 days post infection and incubated at 28°C during treatment.

### Microscopy

For imaging on the fluorescent inverted microscope, embryos were anaesthetised in tricaine as above and mounted in 3% (w/v) methylcellulose in E3 to reduce embryo movement. Images were captured on a Nikon SMZ1500 microscope with NIS-Elements F version 4.00.06 software.

### Statistics

Data analysis was performed as indicated in the figure legends using the GraphPadPrism (version 5) package.

## Results

### Luminescence can be used to monitor zebrafish embryos naturally infected with luciferase-tagged *M. marinum*

To determine if zebrafish embryos infected with bioluminescently labelled *M. marinum* could be monitored using a luminometer, we incubated embryos (n=300) aged 1 and 2 dpf in E3 containing 1x10^7^ cfu ml^-1^ of *M. marinum* tagged either with a modified firefly luciferase (designated *M. marinum* BSG100) or with a modified bacterial luciferase (designated *M. marinum* BSG101). At different time points, we removed the embryos from the *M. marinum*-containing media, washed them in fresh media and measured light. As the firefly luciferase reaction requires luciferin as a substrate, we added this exogenously (30 μg ml^-1^) when embryos were placed into the 96 well plates. Bioluminescence from infected embryos did not rise above background levels until 4 days post infection, with comparable levels of light emitted from BSG100 and BSG101 infected embryos (Fig. 1). Natural infection of embryos with *M. marinum* did not result in any premature deaths when compared to the uninfected group. While embryos infected with *M. marinum* labelled with the firefly luciferase (BSG100) produced more light (median maximum value of 709 relative light units [RLU] (ranging from 328 to 2872)) compared to the bacterial luciferase tagged strain (BSG101) (median maximum value of 446 RLU (ranging from 122 to 796)), the exogenous addition of luciferin increased the time required to carry out the assay and the expense of the technique. We therefore selected bacterial-luciferase tagged *M. marinum* M (BSG101) for further study.

**Figure 1.**
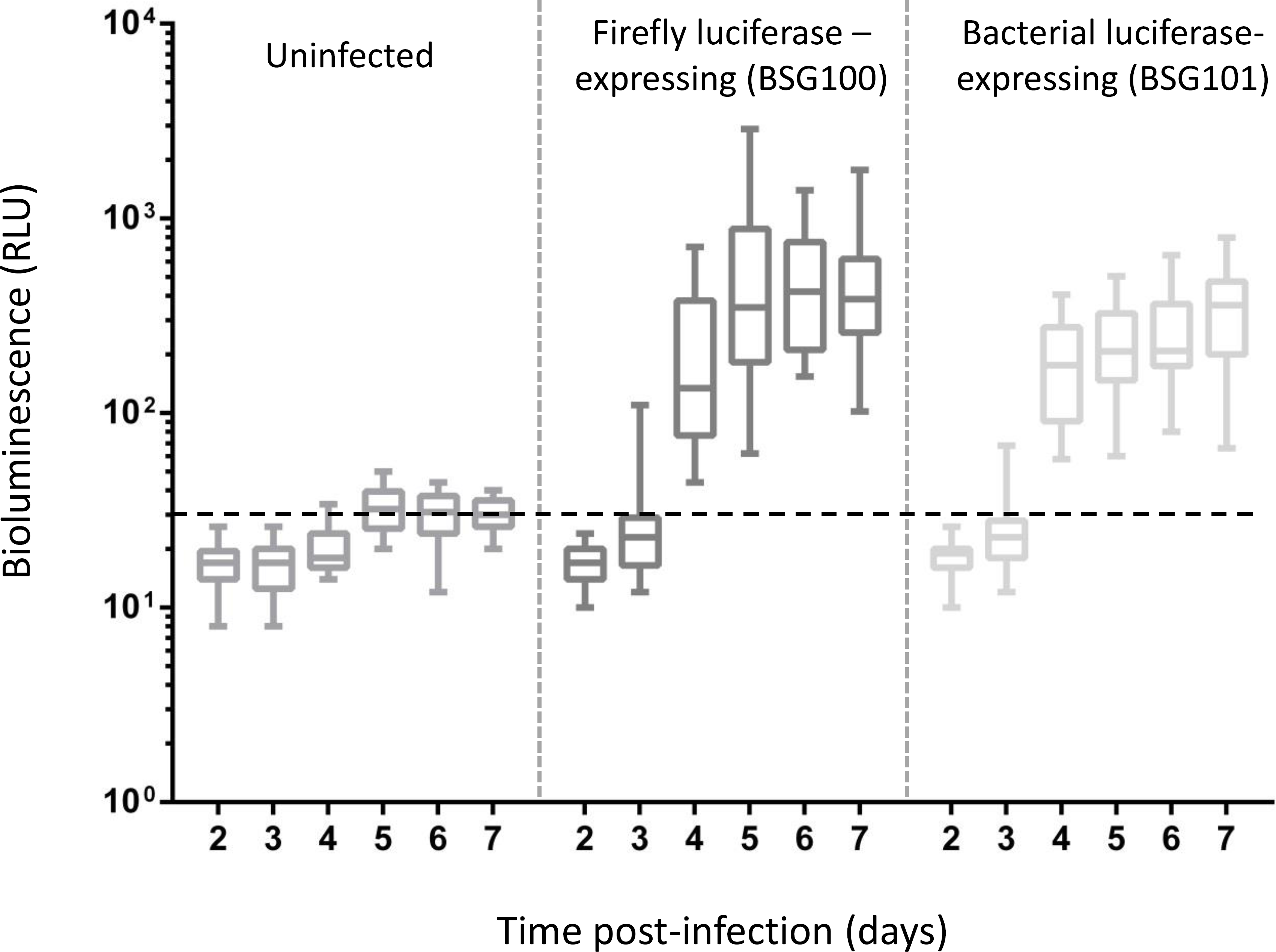
Bioluminescence can be used to monitor zebrafish embryos naturally infected with tagged strains of *M. marinum*. A comparison of the bioluminescence (given as relative light units [RLU]) from zebrafish embryos naturally infected with *M. marinum* M expressing either a red-shifted firefly luciferase (BSG100) or bacterial luciferase (BSG101). Similar light levels were observed with the two constructs while uninfected embryos remained at background levels (indicated by black dashed line). Bioluminescence is presented as box-whisker plots of RLUs from 20-30 embryos measured over a seven day period. The edges of the boxes represent the 25th and 75th quartiles, the solid line represents the median, and the whiskers are the minimum and maximum values. One representative experiment is shown.

### Natural infection results in gill colonisation and transient gut colonisation

We investigated the nature and location of natural *M. marinum* infection of embryos using a strain of *M. marinum* expressing the red fluorescent protein tdTomato^13^ (designated BSG102). We incubated the embryos with BSG102 for 4 days and then determined the location of the infecting bacteria using a fluorescent microscope (Fig. 2). We observed fluorescently tagged bacteria throughout the digestive tract, clustering around the developing gills and lower jaw of the larvae (Fig. 2 A&B). We speculated that those bacteria present within the digestive tract could represent a transient colonisation. These bacteria were seen to be removed following defecation. To test this, we washed infected embryos in fresh media and incubated for a further 24 hours. After this incubation period, we observed that the bacteria that were previously present within the digestive tract had gone, while the bacteria associated with the developing gills were still present (Fig. 2C). We observed that the digestive tract remained clear of detectable bacteria for the majority of the experiment, with some non-transient colonisation appearing approximately 11 days post infection.

**Figure 2.**
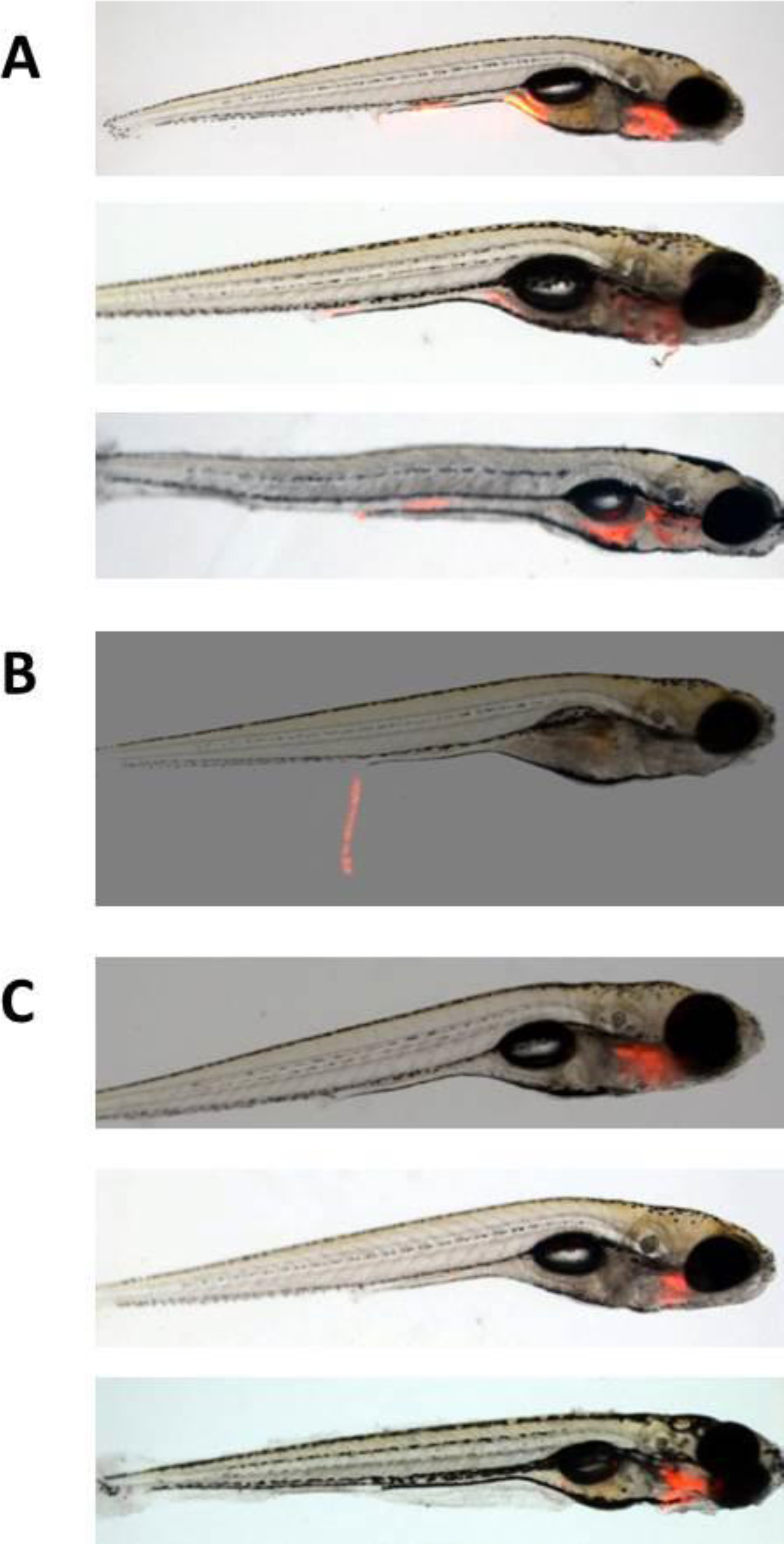
Natural exposure of zebrafish embryos to *M. marinum* results in transient colonisation of the gut and infection of the developing gills and lower jaw. Embryos were exposed to *M. marinum* expressing a red fluorescent reporter (BSG102) and bacterial location identified by fluorescence microscopy. (A) Colonisation of the developing gills and lower jaw after 4 days infection. (B) Transient colonisation of the digestive tract. (C) After 5 days, colonisation is localised to the head region, with the digestive tract no longer colonised. Representative embryos are shown.

### Optimisation of infectious dose and infection protocol

In order to determine the minimum dose for establishing a traceable infection within a single zebrafish embryo, we incubated embryos with concentrations of *M. marinum* BSG101 ranging from 1x10^4^ to 1x10^7^cfu ml^-1^ and followed bioluminescence over 12 days using a luminometer. A dose of 1x10^7^cfu ml^-1^ BSG101 was the only one tested that established an infection that we could reliably detect above background levels using our luminometer (Fig. 3). This dose resulted in approximately 40-60% of the exposed embryos becoming bioluminescent, depending on the experiment (data not shown). When embryos were exposed to a dual bioluminescent/fluorescent tagged strain of *M. marinum* (BSG103) and examined by fluorescence microscopy, we observed that more than 90% of the embryos were infected, but only half of these could be detected by luminometry (data not shown).

**Figure 3.**
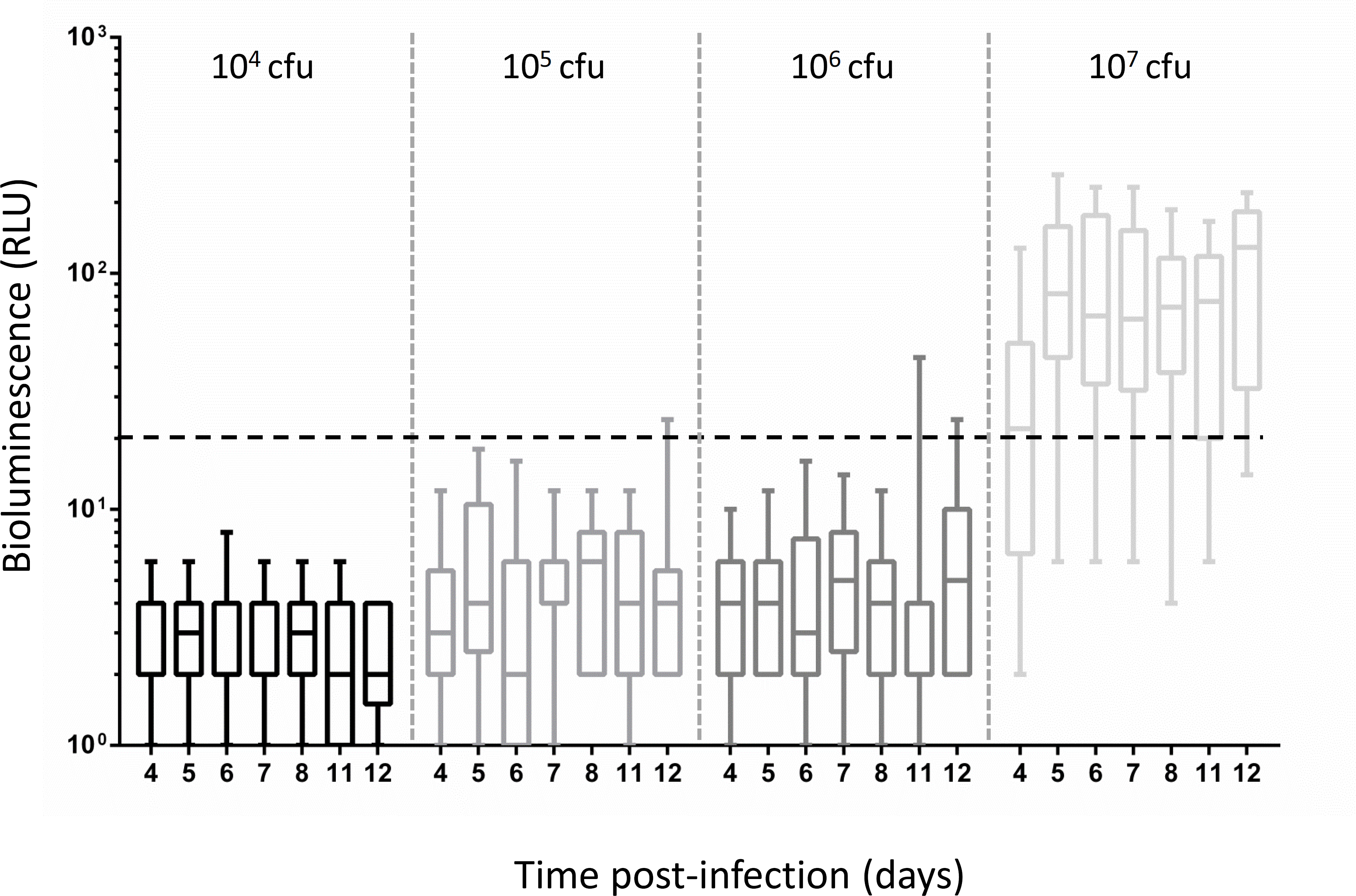
An infectious dose of 10^7^ cfu bioluminescent *M. marinum* per ml of exposed medium is required to produce infected embryos that can be detected by luminometry. A comparison of the bioluminescence (given as relative light units [RLU]) from zebrafish embryos naturally infected with different doses (from 10^4^ to 10^7^ colony forming units [cfu] ml^-1^) of *M. marinum* M expressing bacterial luciferase (BSG101). Black dashed line indicates limits of detection. Bioluminescence is presented as box-whisker plots of RLUs from 20-30 embryos measured over a 12 day period. The edges of the boxes represent the 25th and 75th quartiles, the solid line represents the median, and the whiskers are the minimum and maximum values. One representative experiment is shown.

In an effort to reduce the time required to process embryos for infection, we investigated the effect of manual versus chemical dechorionation (treatment with pronase). We observed that pronase treatment reduced the proportion of infected embryos detectable by luminometry (45% for manual, 7% for pronase) and light levels from infected embryos were also consistently lower (20-150 RLU for manual, 20-76 RLU for pronase). For these reasons, we adopted the following optimised protocol for further experiments: 1) manual dechorionation of embryos 2 dpf, 2) immersion in media containing 1x10^7^cfu ml^-1^ *M. marinum* for 4 days at 28°C, 3) wash in fresh media and incubation for a further 24 hours at 28°C, 4) transfer of individual embryos to the wells of a black 96 well plate using a sterile plastic pasteur pipette for measurement of light and drug intervention studies.

### Drug treatment of naturally infected zebrafish embryos

Zebrafish embryos, infected using the optimised natural infection protocol, were treated with rifampicin and a variety of nitroimidazole-based next generation and experimental anti-mycobacterial drugs (Table 2, Fig. 4). Delamanid was recently approved for clinical use, while Pretomanid is in human Phase III combination trials^39^. The experimental compounds SN30488, SN30527 and SN30982^40^ are analogues of Pretomanid with varying lipophilic side chains, selected for their wide range of potencies against *M. tuberculosis* cultures. For comparison, we treated embryos infected through microinjection into the caudal vein with a subset of the compounds (Fig. 4). We considered all injected embryos as infected, whereas we only selected embryos with observable light emission after natural infection for use in drug intervention studies. For naturally infected embryos we used reduction in light emission as a surrogate measure of anti-mycobacterial activity; for injected embryos, we measured drug efficacy by embryo survival. We also exposed in vitro grown *M. marinum* and *M. tuberculosis* to the same compounds.

**Figure 4.**
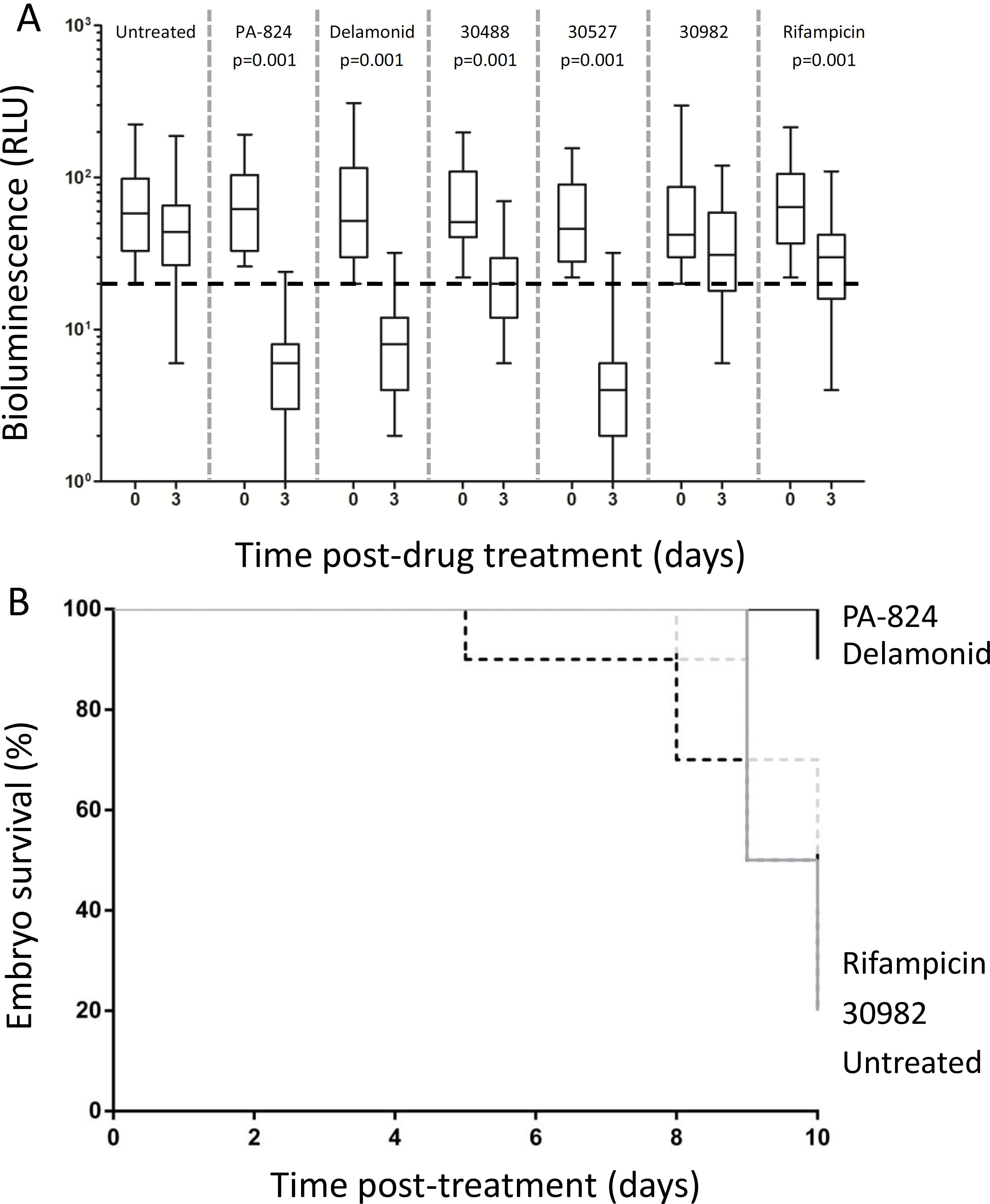
Treatment of *M. marinum* BSG101-infected embryos with Pretomanid, Delamanid, SN30488, SN30527, SN30982 and rifampicin. Drug efficacy was monitored by changes in bioluminescence (given as relative light units [RLU]) from individual naturally infected embryos (n=20-30) immediately prior to and 3 days after treatment (A). Black dashed line indicates limits of detection of luminometer. Data did not pass the D'Agostino & Pearson normality test so before and after treatment groups were compared using the Kruskal Wallis test with Dunn's post-hoc analysis. Those treatments resulting in a significant decrease in bioluminescence are shown. One representative experiment out of 3 is shown. For embryos infected by caudal vein injection (B), after injection embryos were placed directly into media containing compounds and drug efficacy was monitored by changes in survival over ten days. One representative experiment out of 2 is shown.

We observed that at the concentration used, only three of the six treatments tested (Delamonid, SN30527 and rifampicin) retarded the growth of *M. marinum* BSG101 in vitro (as shown by a lack of increase in bioluminescence) (Fig 5A). In contrast, five of the six treatments tested (Pretomanid, Delamanid, SN30488, SN30527 and rifampicin) retarded the growth of *M. tuberculosis* BSG001 in vitro (Fig 5B). Interestingly the same five treatments tested (Pretomanid, Delamanid, SN30488, SN30527 and rifampicin) significantly reduced the bioluminescent signal from naturally infected zebrafish embryos (p<0.001, Kruskal Wallis test with Dunns multiple comparisons) (Fig. 4A). The data is summarised in Table 3. Similarly, we observed a 90% survival rate for BSG101 injected embryos treated with Pretomanid and Delamanid, compared to 20% survival of untreated injected embryos or those treated with SN30982 or rifampicin (Fig. 4B).

**Figure 5.**
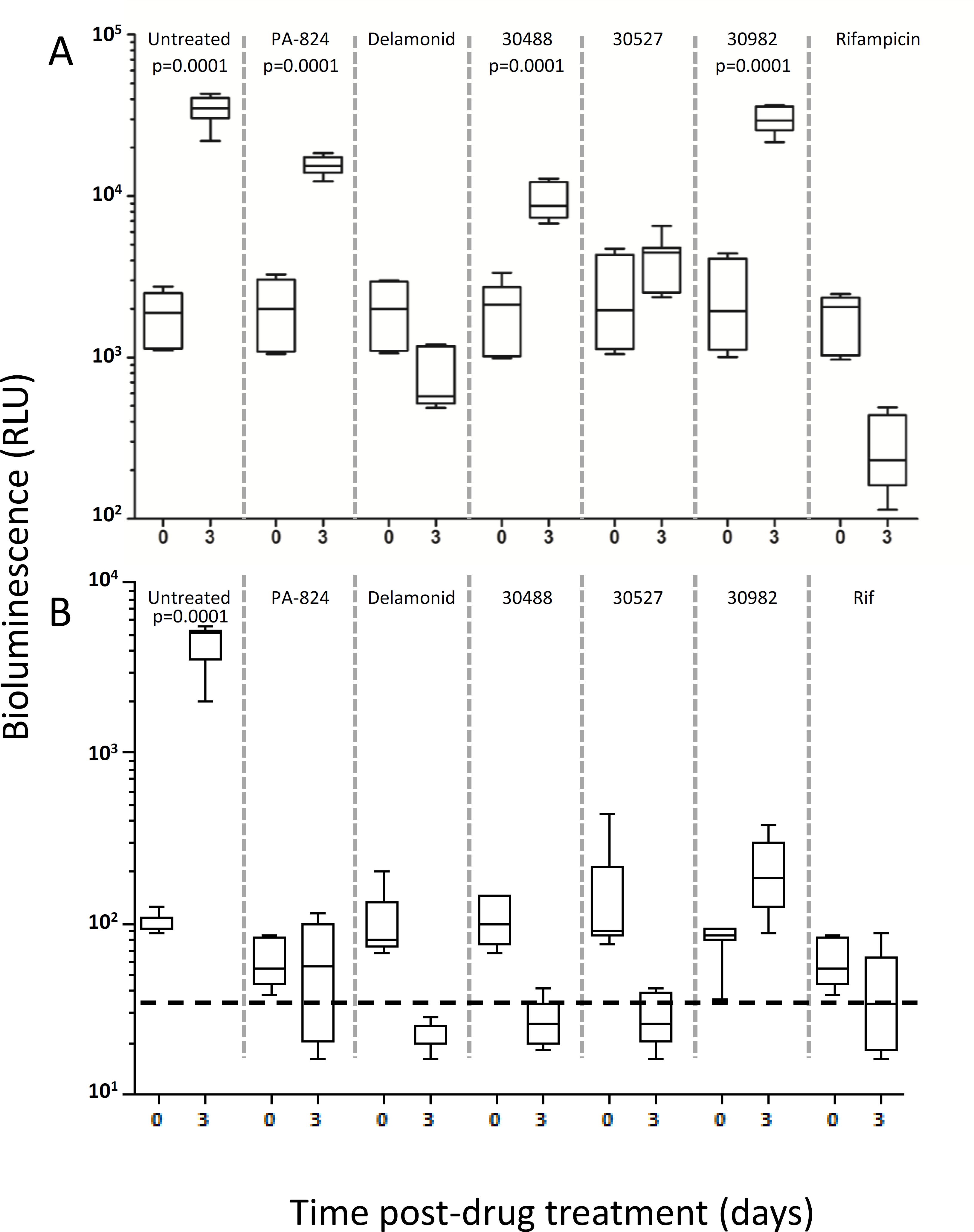
In vitro treatment of *M. marinum* BSG101 and *M. tuberculosis* BSG001 with Pretomanid, Delamanid, SN30488, SN30527, SN30982 and rifampicin. Drug efficacy was monitored by changes in bioluminescence (given as relative light units [RLU]) immediately prior to and 3 days after treatment for *M. marinum* BSG101 (A) or for *M. tuberculosis* BSG001 (B). Data passed the D'Agostino & Pearson normality test so before and after treatment groups were compared using ANOVA with Bonferroni's post-hoc analysis. Those treatments resulting in a significant increase in bioluminescence are shown. Data represents experiments performed on three separate occasions. (Dotted line indicates background level of light detection).

**Table 3.** Comparison of efficacy of treatments across in vitro and in vivo models used in this study.

| Compound ID | In vitro MTB | In vitro MM | Zebrafish MM<br>(natural infection) | Zebrafish MM<br>(caudal vein injection) |
| --- | --- | --- | --- | --- |
| Pretomanid | A | N | A | A |
| Delamanid | A | A | A | A |
| SN30488 | A | N | A | NT |
| SN30527 | A | A | A | NT |
| SN30982 | N | N | N | N |
| Rifampicin | A | A | A | N |

Key: MTB, *M. tuberculosis* BSG001; MM, *M. marinum* BSG101; A, active; N, not active; NT, not tested.

## Discussion

There is a clear and desperate need for new medicines to treat TB. As working with *M. tuberculosis* limits research and preclinical drug development to those laboratories around the world with the resources and facilities to safely handle the bacterium, non-tuberculous mycobacteria such as *M. marinum* are widely used as surrogates^5, 7, 41, 42^. A major drawback of using *M. marinum* for antimycobacterial compound screening is that the in vitro resistance profile of the bacterium is very different to *M. tuberculosis*. This difference can be observed in the drastically different MIC values for the compounds tested against the two organisms in this study (Table 2). Two of the compounds (Pretomanid and SN30488) were not effective against *M. marinum* but were effective against *M. tuberculosis*. This means that screening compounds for activity against *M. marinum* in vitro runs the risk of missing potential anti-TB agents.

Infection of zebrafish, either as adults or as embryos, by injection with *M. marinum* has proved to be a useful model for studying mycobacterial pathogenicity and for drug screening^13, 43^ ^44^. However, current infection protocols require specialised equipment and a high level of operator expertise. We first wanted to establish whether it was possible to infect zebrafish embryos by exposure to *M. marinum* in the media, and to monitor infection dynamics using bioluminescence. We determined that incubation for four days in a petri-dish containing 10^7^ cfu of *M. marinum* per ml of media was sufficient to establish an infection. Indeed, microscopic imaging demonstrated that fluorescent *M. marinum* could be visualised in the developing gills and digestive tract after this time.

The kinetics of natural infection we observed fits with existing understanding of zebrafish larval development; their mouth has been demonstrated to open from 3 dpf and their gut exists as an open-ended tube by 4 dpf^45^. The initial digestive tract colonisation was found to be transitory, although at later stages a non-transitory colonisation was observed. It is unsurprising that the embryos could be infected via bathing, as this should represent one of the ways that zebrafish could become naturally infected in the wild; however without the use of bioluminescence to visualise *M. marinum*, the easy identification of infected embryos would not be possible. We observed that the proportion of embryos which became infected varied depending on the housing conditions. Initially embryos were housed in groups of 50 for infection but this resulted in a low proportion of measurable infections. When we increased the number to 300, the proportion of embryos with a measurable infection rose to approximately 40%. We speculate that this is due to the increased motion of the fish in the media allowing for greater mixing. Another possibility is that *M. marinum* may phenotypically change as it travels through the embryo gut, making it more infectious. Such hyperinfectivity has been reported for *Vibrio cholera* and *Citrobacter rodentium*^46–48^

As several bioluminescent reporter systems exist, we wanted to establish which one would give the best results for this assay. We compared mycobacterial optimised bacterial luciferase (lux) and the codon-optimised red shifted firefly luciferase (RTluc)^49^. The bacterial lux construct contains all the required genes to make both the substrate and the catalytic enzyme to produce light. In the case of the firefly luciferase, the substrate has to be added exogenously. While *M. marinum* tagged with the firefly luciferase did produce higher levels of light, this was not enough to compensate for the increased cost to the assay, both in terms of expense and time as a result of having to add the exogenous substrate. We chose to use *M. marinum* tagged with bacterial lux through-out to produce as streamlined and economic an assay as possible.

We investigated whether the assay could be accelerated by using pronase treatment to dechorionate embryos. While pronase treatment did reduce the time required to prepare embryos, more bacteria were needed to establish an infection compared to the dose needed to infect manually dechorionated embryos. Similarly, the proportion of infected embryos with visible bioluminescence was also reduced. One possible explanation for this difference could be that pronase treatment may result in an increased inflammatory reaction within the embryos, resulting in greater initial killing of the bacteria.

The optimised natural infection assay protocol we have established requires manual dechorination of zebrafish embryos two days post fertilisation, followed by bathing with 1x10^7^ cfu ml^-1^ *M. marinum* for 4 days. Embryos are then removed from the infected media, washed and placed in fresh media and kept for a further overnight to allow for the clearance of transiently colonising bacteria. After a further wash, embryos can be housed within individual wells of a clear bottom, black 96 well plate for measurement of infection dynamics using bioluminescence. We investigated whether the optimised natural infection assay could be applied to the testing of anti-mycobacterial compounds. Four of the six compounds we tested reduced the luminescence from infected embryos to below background levels. When the same compounds were tested in the caudal vein-injection model, which uses embryo survival as an indicator of bacterial clearance, three of the four compounds were also identified as being effective.

Overall this study has demonstrated that it is possible to carry out high throughput in vivo drug screening in the zebrafish model using embryos naturally infected with bioluminescent *M. marinum* M. Natural infection is quicker than injection and requires less expertise. Interestingly, not all of the injected embryos had detectable light levels and with the fluorescently tagged *M. marinum* up to 90% of the naturally infected embryos had visible signs of infection (data not shown). Embryos can be screened in 96 well plates and drug efficacy rapidly identified over the course of 10 days. Through the use of a luminometer with a plate stacker this process can be semi-automated to reduce the hands on time. While this is moderately slower than previously reported automated robotic systems, it is also a fraction of the cost. The result is an assay that can be carried out by a wide variety of laboratories for minimal cost and without high levels of zebrafish expertise. Widening participation in TB research and pre-clinical drug discovery in this way should accelerate the progress towards new and better treatments for TB and other neglected mycobacterial infections.

## Acknowledgements

The authors would like to thank Professor Lalita Ramakrishnan for the kind gift of plasmid pTEC27.

## Transparency statement

No conflicts of interest to declare.

